# Interleukin (IL)-17/IL-36 axis participates to the crosstalk between endothelial cells and keratinocytes during inflammatory skin responses

**DOI:** 10.1101/767400

**Authors:** Cristina M. Failla, Lorena Capriotti, Claudia Scarponi, Laura Mercurio, Francesco Facchiano, Martina Morelli, Stefania Rossi, Gianluca Pagnanelli, Cristina Albanesi, Andrea Cavani, Stefania Madonna

**Affiliations:** Laboratory of Experimental Immunology, IDI-IRCCS, Rome, Italy; Department of Oncology and Molecular Medicine, Istituto Superiore di Sanità (ISS), Rome, Italy; 1st Dermatology Division, IDI-IRCCS, Rome, Italy; National Institute for Health, Migration and Poverty (NIHMP), Rome, Italy

**Author notes:** Department of Medical-Surgical Sciences and Biotechnologies, Sapienza University of Rome, Italy (Lorena Capriotti). Corresponding author: Cristina M. Failla. These authors contributed equally to the work (Andrea Cavani, Stefania Madonna).

## Abstract

In inflammatory skin conditions, such as psoriasis, vascular enlargement is associated with endothelial cell proliferation, release of cytokines and adhesion molecule expression. Interleukin (IL)-17A is a pro-inflammatory cytokine mainly secreted by T helper-17 cells that is critically involved in psoriasis pathogenesis. IL-36α, IL-36β and IL-36γ are also inflammatory cytokines up-regulated in psoriasis and induced by various stimuli, including IL-17A. In this study, we found that human keratinocytes are the main source of IL-36, in particular of IL-36γ. This cytokine was strongly induced by IL-17A and efficiently activated human dermal microvascular endothelial cells (HDMECs), which expressed both IL-17 and IL-36 receptors, by inducing a molecular signaling, such as phosphorylation of ERK1/2 and NF-κB P65 subunit. We highlighted the intense IL-17A- and IL-36γ-dependent interplay between keratinocytes and HDMECs, likely active in the psoriatic lesions and leading to the establishment of a cytokine network responsible for the development and maintenance of the inflamed state. On HDMECs, IL-17A or IL-36γ showed a synergic activity with TNF-α, potently inducing inflammatory cytokine/chemokine release and ICAM-1 expression. We also investigated the involvement of IL-36γ and VEGF-A, substantially reduced in lesional skin of psoriatic patients pharmacologically treated with the anti-IL-17A antibody Secukinumab. Importantly, keratinocyte-derived IL-36γ represented an additional pro-angiogenic mediator of IL-17A. We observed that keratinocyte-derived VEGF-A influenced proliferation but not reduced inflammatory responses of HDMECs. On the other hand, inhibition of IL-36γ released by IL-17A-treated keratinocytes impaired ICAM-1 expression in HDMECs. Taken together, our data demonstrated that IL-17A and IL-36γ are highly involved in endothelial cells/keratinocytes crosstalk in inflammatory skin conditions.

## Introduction

Blood and lymphatic vessels have a major role in skin inflammation [1]. In chronic inflammatory disorders, such as psoriasis, vascular enlargement is associated to vessel hyper-permeability and endothelial cell (EC) proliferation. Vessel morphological changes are evident well before the development of epidermal hyperplasia, even if most pro-angiogenic factors are produced by epidermal keratinocytes themselves [2]. Besides, activated endothelium expresses adhesion molecules and secretes cytokines and chemokines that support leukocyte extravasation and migration into the skin, thus contributing to disease pathogenesis [3]. Under inflammatory conditions, MHC class II^+^ ECs have been also involved in the selective amplification of interleukin (IL)-17-producing CD4^+^ T helper (Th) lymphocytes [4,5]. IL-17 cytokines, in particular IL-17A, are potent proinflammatory cytokines secreted by Th-17 cells and by additional adaptive and innate lymphocytes as well as neutrophils and mast cells [6]. The IL-17 family comprises six members that exert their functions as homodimers with the exception of IL-17A and IL-17F that can form heterodimers. In a similar way, IL-17 cytokines signal via heterodimeric receptors (IL-17R) and IL-17A, IL-17F or IL-17A/IL-17F heterodimers bind to the same receptor composed of IL-17RA and a IL-17RC subunits. IL-17RA is ubiquitously expressed in epithelial, hematopoietic cells, fibroblasts and osteoblasts, as well as ECs [7]. However, IL-17 family involvement in EC biological responses is still a controversial issue, especially in inflammatory conditions. Tumors expressing IL-17A show a high vascular density, and IL-17A elicits neovascularization in a rat cornea assay [8]. Some authors reported that IL-17A does not directly affect endothelial cell proliferation *in vitro* [8] but significantly enhances proliferation induced by other angiogenic cytokines such as vascular endothelial growth factor (VEGF)-A [9]. Moreover, IL-17A induces EC migration and tubular structure formation *in vitro* [8]. Other studies reported a direct role of IL-17A in vessel growth *in vitro* and *in vivo*, through activation of both IL-17RA and IL-17RC [10]. Furthermore, Liu *et al*. reported that IL-17A effects on vascular inflammation were not mediated by ECs but rather by pericytes [11].

On ECs and other cell types, most of the IL-17A-induced inflammation depends on its capability to act synergistically with other stimuli. IL-17A and IL-6 together induce ICAM-1 up-regulation in ECs, enhancing monocyte adhesion to vessels [12]. In the case of tumor necrosis factor (TNF)-α, IL-17A stabilizes the mRNA of TNF-activated genes leading to a signal amplification [13]. IL-17 and TNF-α synergistically stimulate cytokine expression in human melanocytes and ECs [14,15]. In human dermal microvascular ECs (HDMECs), IL-17A cooperates with TNF-α in the induction of CSF1/G-CSF and CXCL1/GRO-α [14], whereas in brain ECs IL-17A alone stimulates the release of CCL2/MCP-1 and CXCL1/GRO-α [16]. On an immortalized endothelial cell line, IL-17A was able to stimulate the release of CXCL1/GRO-α, CSF2/GM-CSF and CXCL8/IL-8 [17]. Therefore, literature data are highly variable, considering the different origin of the utilized ECs and the diverse culture conditions.

IL-17A is critically involved in the pathogenesis of psoriasis and several drugs targeting the IL-17A pathway have been developed and are currently used in the clinical practice. IL-17A affects, in particular, keratinocyte immune function, by inducing the release of antimicrobial peptides and chemokines, such as CXCL8/IL-8 and CXCL1/GRO-α, responsible for the accumulation of neutrophils in the early phase of psoriasis inflammation [18]. Among the factors induced by IL-17A, together with TNF-α, there are the IL-36 cytokines that, in turn, augment Th-17 functions, revealing the existence of a feedback loop able to amplify the IL-17 inflammatory signals [19]. IL-36 cytokines belong to the IL-1 family and are highly present in psoriasis, being produced by keratinocytes, macrophages and dendritic cells [20]. IL-36α, β and γ initiate a signal cascade that starts with binding to their IL-1Rrp2 receptor and leads to up-regulation of proinflammatory cytokines including IL-6 and CXCL-8/IL-8 [21]. In the psoriasis context, IL-36 cytokines, together with IL-17A, impair keratinocyte differentiation by inducing a proinflammatory skin phenotype [22]. Importantly, IL-36 family impacts on immune response initiation by acting on dendritic and Langerhans cells, recruiting neutrophils, and promoting CD4+ T cell proliferation [23,24]. HDMECs also express IL-36 receptor and respond to IL-36γ stimulation by up-regulating adhesion molecule expression and augmenting chemokine secretion [25]. Less is known about a possible role of IL-36 in the crosstalk between ECs and epidermal keratinocytes.

In this paper, we investigated direct and indirect effects of both IL-17A and IL-36γ on HDMECs, underlining the importance of these cytokines in the crosstalk between keratinocytes and ECs during skin inflammatory processes.

## Materials and methods

### Cell culture

Human keratinocytes were obtained from skin biopsies of healthy donors as previously described [26]. Experiments were carried out on secondary and tertiary cultures and repeated at least three times on different strains. HDMECs were isolated from foreskin of donors as previously described [27] and used at passage 4 to 8. A pool of HDMECs derived from 4 different healthy donors was used.

### Western blotting

HDMECs were starved in endothelial basal medium (EBM, Basel Switzerland) supplemented with 2% fetal bovine serum for 6 hours and then left untreated or treated with 10 ng/ml IL-17A or 20 ng/ml IL-36γ alone or with the addition of 10 ng/ml TNF-α (R&D Systems, Minneapolis, MN, USA) for 24 hours. Cells were lysed in RIPA buffer [50 mM Tris-HCl (pH 7.4), 150 mM NaCl, 1 mM EGTA, 1% NP-40, 0.25% Na deoxycholate, 0.1% SDS] and 30 μg of the total protein lysate were loaded on a 10% SDS-polyacrylamide gel, transferred to nitrocellulose (Hybond-ECL, GE Bioscience, Chalfont St.Giles, UK) and incubated for 1 hour in Western blocking reagent (Roche Applied Science, Basel, Switzerland). Primary antibodies (anti-human IL-17RA antibody, Cell Signaling Technology, Danvers, MA, USA; anti-human IL-1Rrp2 antibody, Santa Cruz Biotechnologies, Santa Cruz, USA) were used diluted 1:1000 and applied for 18 hours followed by the appropriate horseradish peroxidase-coupled secondary antibody (GE Bioscience).

Blots were re-probed with anti-β-actin antibody (diluted 1:4000, Santa Cruz Biotechnology, Santa Cruz, CA, USA) and stained with Coomassie blue as loading controls as previously described [28]. Detection was performed using the ECL plus detection system (GE Bioscience). The relative intensity of signals was quantified using a GS-710 densitometer (Bio-Rad Laboratories, Hercules, CA, USA). For signaling studies, HDMECs were treated or not with IL-17A (50 ng/ml) or IL-36γ (20 ng/ml) for different times and lysed as aforementioned. Western blotting analyses were performed by using the following primary antibodies: mouse anti-phosphorylated (p)STAT3 (Cell Signaling Technology); mouse anti-pERK1/2 (Santa Cruz Biotechnologies); anti-pP65 (Cell Signaling Technology); followed by the appropriate horseradish peroxidase-coupled secondary antibody (GE Bioscience). Blots were re-probed with anti-STAT3, anti-ERK1/2 and anti-P65 antibodies against the not-phosphorylated protein forms (all purchased from Santa Cruz Biotechnologies). The relative intensity of signals was quantified using a GS-710 densitometer (BioRad).

### ELISA assay

HDMECs were seeded in 12-well plates, starved in EBM supplemented with 2% fetal bovine serum for 6 hours and then untreated or treated with 10 ng/ml IL-17A, 10 ng/ml TNF-α, 20 ng/ml IL-36γ or a combination of IL-17A or IL-36γ plus TNF-α for 24 hours. Keratinocytes were seeded in 12-well plates and stimulated with 10 ng/ml IL-17 or with a combination of 200 U/ml interferon (IFN)-γ and 50 ng/ml TNF-α or with the three cytokines together for 24 hours in keratinocyte basal medium (KBM, Lonza). Supernatants were collected, cleared by centrifugation, and attached cells were detached and counted by trypan blue colorimetric assay. Duo Set ELISA kits (R&D Systems) were used for IL36α, IL-36β, IL-36γ, VEGF-A, IL-6, CSF3/G-CSF, CXCL10/IP10 and IL-8. For CCL2/MCP1 and CCL5/RANTES detection BD ELISA kits (OptEIA Set, BD Biosciences, San Diego, CA) were used. Results were normalized to the total number of cells in each sample and were expressed as pg or ng/10^6^ cells. Triplicate wells were used for each condition and experiments were repeated at least three times with comparable results.

### Cell proliferation

HDMECs were plated in a 6 multi-well at the concentration of 1 x 10^5^ cells/ml. At 40% confluence, cells were starved in EBM supplemented with 2% fetal bovine serum for 4 hours and treated with: i) IL-17A (10 and 50 ng/ml), alone or in combination with 10 ng/ml of TNF-α; ii) IL-36γ (50 ng/ml), alone or in combination with either 10 ng/ml of TNF-α or 10 ng/ml IL-17A, in EBM plus 2% fetal bovine serum; iii) the endothelial growth medium (EGM, Lonza) or iv) untreated. The number of viable cells was determined by a trypan blue exclusion test after 24, 48 and 72 hours at the end of stimulation. In selected experiments, HDMECs were treated with keratinocyte-conditioned medium. Briefly, keratinocytes were seeded at a concentration of 0.4 x 10^5^ cells/ml in 6-well plate and, reached about 70% confluence, cells were stimulated for 3 hours with IL-17A (50 ng/ml), alone or in combination with TNF-α (50 ng/ml) in KBM. Medium with stimuli was removed and basal medium was added for 48 hours. Next, HDMECs were seeded in 12-well plates in EGM and 1 day after cells were starved and treated with conditioned medium of keratinocytes, in the presence or not of IL-36RA (R&D Systems, 100 ng/ml) or Sunitinib (Pfizer S.r.l., New York, NY, 400 nM). After 24 hours of stimulation, the number of viable cells was determined by a trypan blue exclusion test. At least three independent experiments were performed.

### FACS analysis

Cells were treated as described in the “Cell proliferation” section and HDMEC membrane expression of ICAM-1 was evaluated using allophycocyanin (APC)-conjugated anti-CD54 (ICAM-1) monoclonal Ab (clone 84H10; Immunotech, Marseille, France). VCAM-1 expression and E-selectin expression were detected by using APC-conjugated monoclonal antibodies anti-CD106 (VCAM-1, clone 51-10C9) and anti-CD62E (E-selectin, clone 68-5H11, BD Biosciences) respectively. Cells were analyzed by a FACScan equipped with Cell Quest software (Becton Dickinson, Mountain View, CA, USA). At least three independent experiments were performed.

### Cytokine analysis

HDMECs were starved in EBM supplemented with 2% fetal bovine serum for 6 hours and then untreated or treated with 10 ng/ml IL-17 or TNF-α or both in EBM plus 2% fetal bovine serum for 24 hours. In a different experiment, HDMECs were treated with 20 ng/ml IL-36γ or 10 ng/ml TNF-α or both in EBM plus 2% fetal bovine serum for 24 hours. Conditioned medium was collected and analyzed by means of xMAP technology using a X200 Luminex platform (Bio-Plex) equipped with a magnetic workstation. Panels used were PRO Human Cytokine 27-PLEX (FGF basic, Eotaxin, G-CSF, GM-CSF, IFN-γ, IL-1β, IL-1ra, IL-2, IL-4, IL-5, IL-6, IL-7, IL-8, IL-9, IL-10, IL-12 (p70), IL-13, IL-15, IL-17A, IP-10, MCP-1 (MCAF), MIP-1α, MIP-1β, PDGF-BB, RANTES, TNF-α, VEGF-A) and PRO Human Chemokine 40-PLEX (6Ckine/CCL21, BCA-1/CXCL13, CTACK/CCL27, ENA-78/ CXCL5, Eotaxin/CCL11, Eotaxin-2/CCL24, Eotaxin-3/CC L26, Fractalkine/CX3CL1, GCP-2/CXCL6, GM-CSF, Groα/CXCL1, Gro-β/CXCL2, I-309/CCL1, IFN-γ, IL-1β, IL-2, IL-4, IL-6, IL-8/CXCL8, IL-10, IL-16, IP-10/CXCL10, ITAC/CXCL11, MCP-1/CCL2, MCP-2/CCL8, MCP-3/CC L7, MCP-4/CCL13, MDC/CCL22, MIF, MIG/CXCL9, MIP1α/CCL3, MIP-1δ/CCL15, MIP-3α/CCL20, MIP-3β/CCL 19, MPIF-1/CCL23, SCYB16/CXCL16, SDF-1α + β/CXCL12, TARC/CCL17, TECK/CCL25, TNF-α) (BioRad). Data were analyzed using Bio-Plex Software Manager 6.1. Triplicate wells were used for each condition. Coefficient of variation (CV) of measurements of the whole panel was always lower than 10%.

### Immunohistochemistry

Six-mm punch skin biopsies of three patients with mild-to-severe chronic plaque psoriasis undergoing to pharmacological treatment with the anti-IL-17A antibody Secukinumab (Cosentyx, Novartis Farma S.p.A., subcutaneous injection of 300 mg, once a week after an induction phase) were analyzed by immunohistochemistry. For each patient, biopsies were taken before treatment from lesional (LS) and non-lesional (NLS) skin areas, as well as at LS sites after 8-week treatment. Patients received information and gave their consent to participate to the study. The latter was approved by IDI-IRCCS Ethical Committee (IDI-IMM-IL36pso, registration no. 475/1) and performed in accordance with the Helsinki Declaration. Skin samples were fixed in 10% formalin and embedded in paraffin. Five-μm sections were dewaxed and rehydrated. After quenching endogenous peroxidase, achieving antigen retrieval and blocking nonspecific binding sites, sections were incubated with the anti-VEGF-A mouse monoclonal antibody (Beckton Dickinson) or anti-IL-36γ (AbCam, Cambridge, UK) at a concentration of 5 μg/ml. Secondary biotinylated mAbs and staining kits were obtained from Vector Laboratories. Immunoreactivity was visualized with peroxidase reaction using 3-amino-9-ethylcarbazole (AEC, Vector Laboratories, Burlingame, CA) in H2O2 and specimen counterstained with hematoxylin. As a negative control, primary Abs were omitted or replaced with an irrelevant isotype-matched mAb. Stained sections were analyzed with the AxioCam digital camera attached to the Axioplan 2 microscope (Carl Zeiss AG, Oberkochen, Germany).

### Statistical analysis

Statistical analysis was done using a two-tailed paired Student’s t-test. Statistically significant differences were defined as p≤0.05.

## Results

In order to investigate the role played by IL-17A and IL-36 cytokines in the activation of skin EC, expression of IL-17 and IL-36 receptors by HDMECs was firstly analyzed, together with possible modulation of this expression by inflammatory cytokines present in the psoriasis microenvironment, such as TNF-α. As shown in Fig 1A, Western blotting analysis confirmed the previously reported expression of IL-17 (IL-17RA) and IL-36 (IL-1Rrp2) receptor subunits in HDMECs (Fig. 1A) [11,14]. Densitometric analysis (not shown) indicated that IL-17RA and IL-1Rrp2 expression was not significantly influenced by treatment with IL-17A and IL-36γ, alone or in combination with TNF-α (Fig.1A).

**Fig 1.**
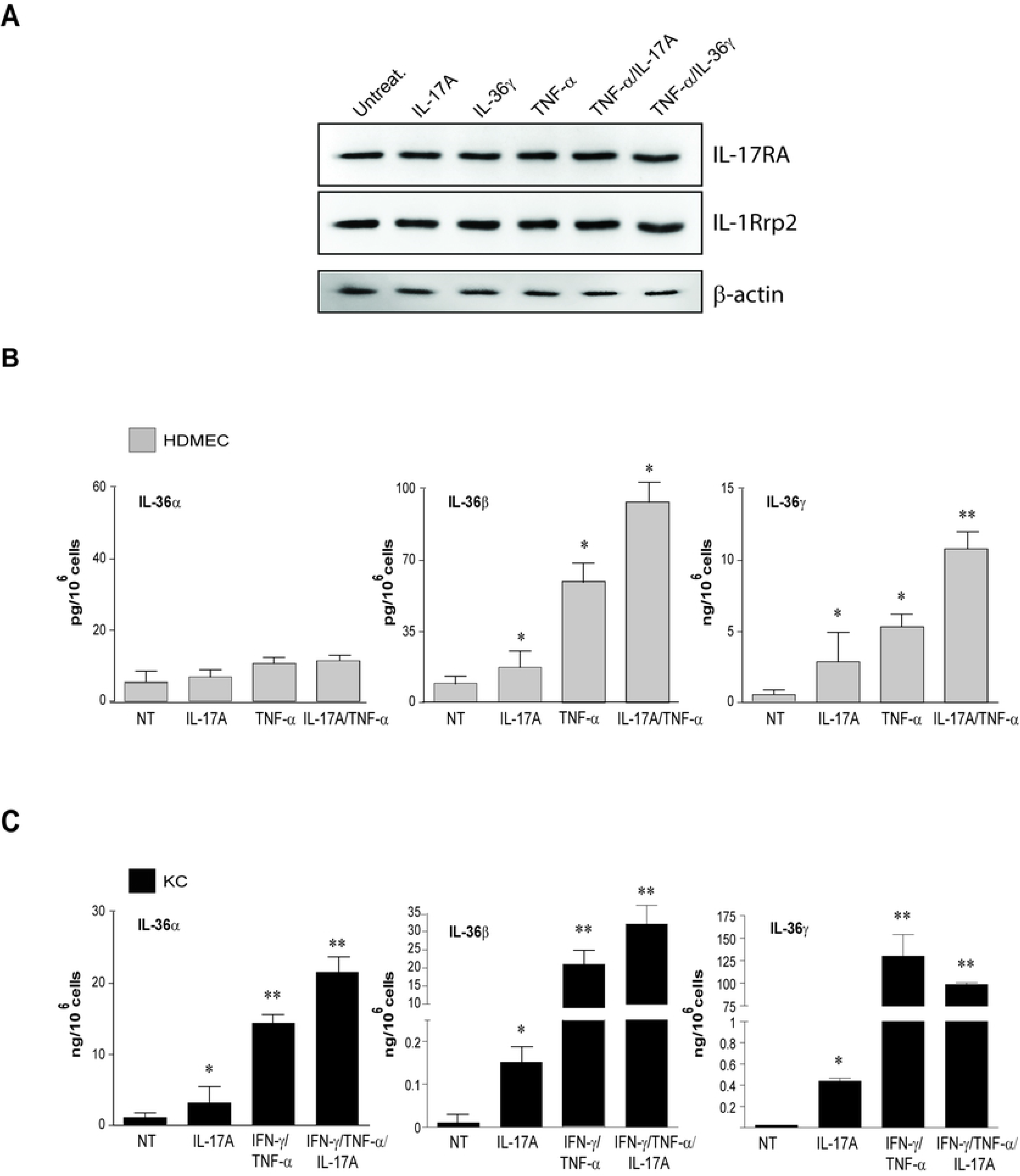
HDMECs express IL-17 and IL-36 receptors and secrete IL-36α, β and γ cytokines. **A.** Protein extracts were obtained from HDMECs stimulated for 24 hours with either IL-17A, IL-36γ, TNF-α or combination of the cytokines (TNF-α+IL-17A; TNF-α+IL-36γ). Western blotting analysis was performed to detect IL-17RA and IL-1Rrp2 expression; β-actin expression was used as a loading control. One representative Western blotting of three different performed is shown. **B-C.** Release of IL-36α, β and γ was analyzed by ELISA in the supernatants obtained by HDMECs (B) and keratinocytes (KC) (C) after a 24 hour cytokine stimulation. All data shown are the mean of pg or ng/10^6^ cells ± SD from three independent experiments; *p ≤ 0.05 and \*\**p* ≤ 0.01 compared with untreated cells.

Differently from IL-17 isoforms that are mainly produced by leukocytes, IL-36 cytokines are expressed by both HDMECs and keratinocytes [28]. As shown in Fig 1B, HDMECs produced substantial amount of two of the three IL-36 isoforms with their release augmenting upon IL-17A or TNF-α stimulation and even more with their combination (Fig. 1B). However, IL-36 levels were lower than those secreted by human keratinocytes (Fig. 1C). Consistently with our recent findings [28], among all the IL-36 isoforms produced by keratinocytes, IL-36γ amount was significantly higher (Fig. 1C) and for these reasons the following experiments were performed with IL-36γ only.

It is known that binding of IL-17A to its receptor induces the intracellular pathways mediated by NF-κB and p38/MAPK in several cell types [29]. As shown in Fig 2A, we found that in HDMECs IL-17A induced the phosphorylation of the transcription factor STAT3 at late time-points of stimulation (6-18 hours). Additionally, IL-17A had a dual effect on ERK1/2 phosphorylation (Fig 2A). In particular, phosphorylation of ERK1/2 was up-regulated rapidly after a 5-min treatment (early activation) with IL-17A and gradually declined after 15 min, although its levels remained higher than those observed in untreated cells. After 6 hours (late activation), ERK1/2 phosphorylation returned to be high and decreased thereafter (Fig 2A). Finally, consistently with other reports, IL-17A induced in HDMECs the early phosphorylation of P65, a transcription factor of the NF-κB complex [29]. Differently from IL-17A, IL-36γ did not activate the phosphorylation of STAT3, whereas it strongly induced the phosphorylation of P65 and, at a lower extent, that of ERK1/2 (Fig 2B).

**Fig 2.**
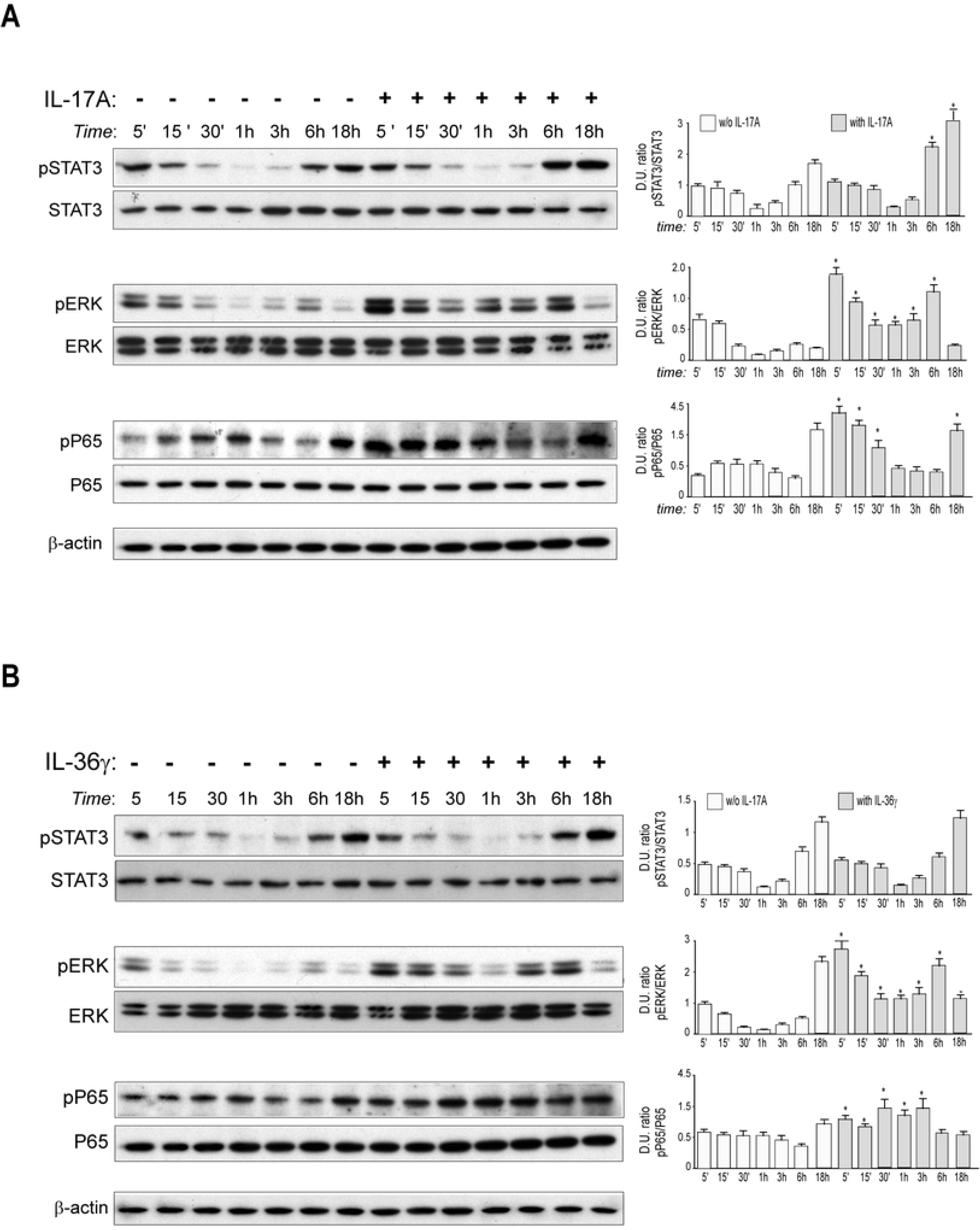
Intracellular signaling induced by IL-17A and IL-36γ in HDMECs. **A and B.** Protein extracts were obtained from HDMECs treated with IL-17A (A) or IL-36γ (B) for the indicated time points and subjected to Western blotting analysis to detect STAT3, ERK1/2 and P65 phosphorylation. Filters were re-probed with anti-STAT3, -ERK1/2 and -P65 antibodies, whereas β-actin levels were detected as a loading control. One representative experiment out of three performed is shown. Graphs in (A) and (B) represent densitometric analyses of the indicated proteins obtained in a representative Western blotting. Data are expressed as mean± SD of the ratio of the Densitometric Units (D.U.) between the indicated phosphorylated/unphosphorylated proteins; *p*\*≤0.05.

We next investigated whether IL-17A or IL36γ could directly influence HDMEC proliferation. As shown in Fig 3A (left panel), HDMEC proliferation was significantly promoted by IL-17A in a dose-response manner at 48 and 72 hours of treatment, as compared to cultures grown in basal medium (EBM). Moreover, when IL-17A was given in combination with TNF-α, it partially reverted the anti-proliferative effect of TNF-α within the 48 hours, at both 10 ng/ml and 50 ng/ml concentrations (Fig. 3A, right panel). At 72 hours of treatment, despite of the reduction of the number of viable cells, the presence of IL-17A at both concentrations contributed to the survival of a higher number of HDMECs, compared to HDMECs treated with TNF-α alone (Fig 3A, right panel). Similarly, IL-36γ significantly promoted HDMEC proliferation even if less efficiently than IL-17A, and the association of the two cytokines did not further enhance cell proliferation (Fig 3B, left panel). However, differently from IL-17A, IL-36γ had a limited potential in protecting cells from the anti-proliferative effect of TNF-α (Fig 3B, right panel).

**Fig 3.**
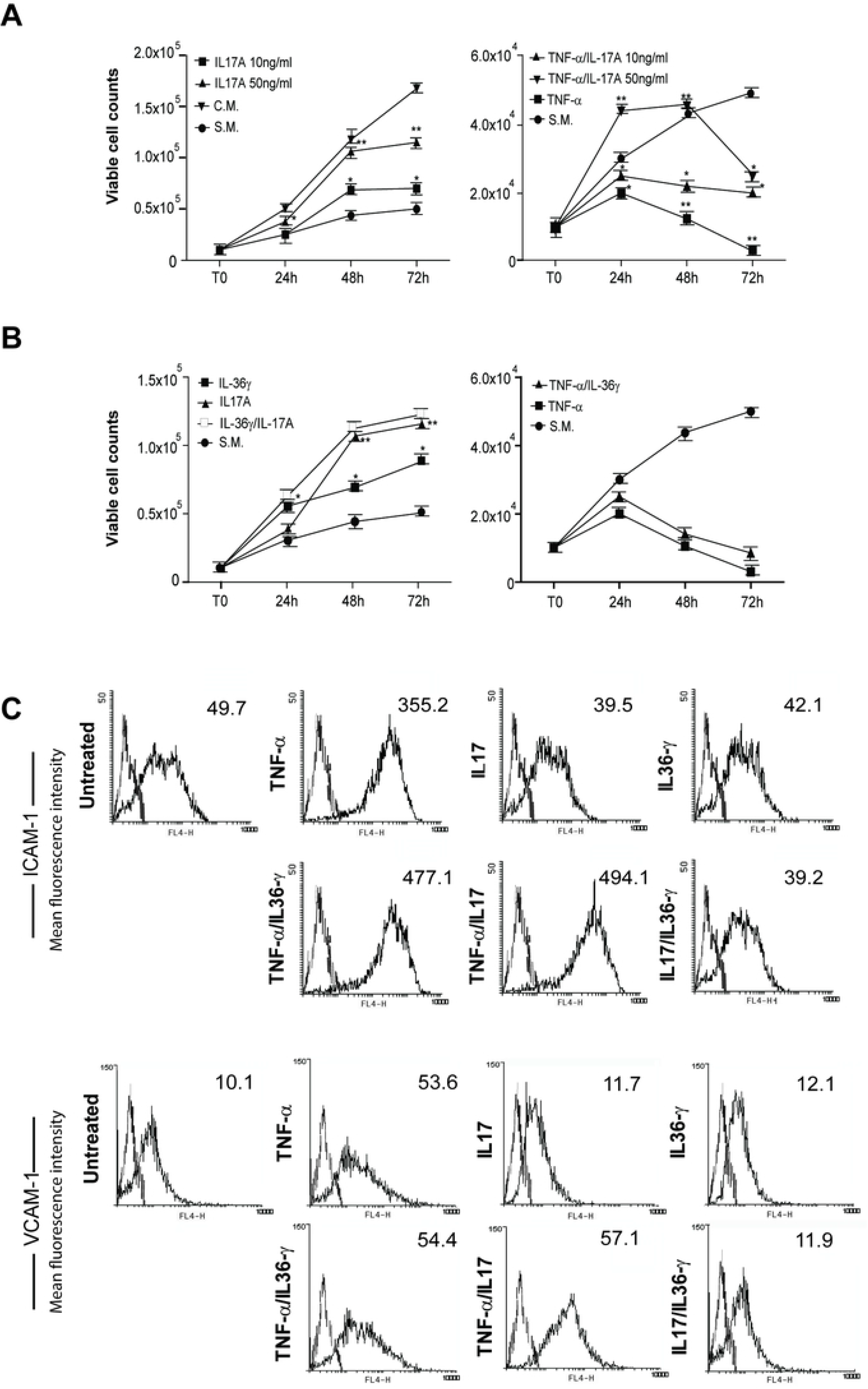
Both IL-17A and IL36γ induced HDMEC proliferation and synergize with TNF-α to induce ICAM-1 expression. **A.** HDMEC cultures were grown in EGM as a complete medium (C.M.) or in EBM as a starvation medium (S.M.) in the presence or absence of 10 or 50 ng/ml IL-17A, administered alone or in combination with TNF-α (10 ng/ml) for the indicated time points. **B.** HDMECs were treated with IL-36γ, alone or in combination with IL-17A or TNF-α in S.M. or untreated. Data are shown as mean values of viable cell counts (trypan blue exclusion test) obtained from three independent experiments ± SD. \**p*≤0.05, ** *p*≤0.01. **C.** ICAM-1 and VCAM-1 expression was evaluated by flow cytometry analysis on HDMECs stimulated for 48 hours with IL-36γ (50 ng/ml), IL-17A (50 ng/ml), TNF-α (10 ng/ml) alone or with combinations of these cytokines, and expressed as mean of the fluorescence intensity (Δmfi). Data shown represent one out of three independent experiments.

In inflammatory conditions, ECs up-regulate membrane receptors that are fundamental for leukocyte adhesion and extravasation from the blood flow into the inflamed tissue [30]. To study the expression of adhesion molecules on the HDMEC membrane following treatment with IL-17A or IL-36γ, flow cytometry analysis was performed on HDMECs treated with: i) IL-17A, alone or in combination with TNF-α; ii) IL-36γ, alone or in combination with TNF-α; iii) IL-17A in combination with IL-36γ; for 48 hours. As shown in Fig 3C, treatment of HDMECs with IL-17A or IL-36γ alone did not affect membrane expression of ICAM-1. In a similar way, HDMEC treatment with IL-36γ in combination with IL-17A did not influence membrane expression of ICAM-1. On the other hand, both IL-17A and IL-36γ significantly synergized with TNF-α in the induction of ICAM-1. No modulation of expression of either VCAM-1 (Fig 3C) or E-selectin (data not shown) could be observed after IL-17A or IL-36γ treatment.

To investigate the effects of IL-17A and IL-36γ treatment on cytokine and chemokine secretion by HDMECs, we used Bio-Plex Pro™ assays in which several inflammatory molecules could be simultaneously analyzed. As a comparison, HDMECs were treated with TNF-α alone or in combination with either IL-17A or IL-36γ. As shown in Table 1, IL-6, CXCL8, G-CSF CXCL10 and CCL2 showed a significant up-regulation upon cell treatment with IL-17 alone, even if to a less extent compared to the secretion due to the TNF-α treatment. Secretion of these five cytokines, together with IL-1RA, was significantly augmented upon stimulation with both IL-17A and TNF-α, with an additive effect in respect to treatment with TNF-α alone.

**Table 1.** Bio-Plex 27-Plex cytokine analyses of supernatants from HDMECs treated with IL-17A alone or in combination with TNF-α.

| Molecules | Untreated | | TNF-α | | IL-17A | | TNF-α + IL-17A | |
| --- | --- | --- | --- | --- | --- | --- | --- | --- |
|  | Mean | SD | Mean | SD | Mean | SD | Mean | SD |
| IL-1RA | 5.4 | 0.2 | 121.0<sup>§</sup> | 7.1 | 18.0 | 1.5 | 195.9<sup>#</sup> | 4.8 |
| IL-6 | 72.5 | 13.0 | 5579.0<sup>§</sup> | 506.0 | 1287.1<sup>§</sup> | 147.1 | 13355.3<sup>#</sup> | 475.0 |
| CXCL8 | 157.0 | 6.4 | 2512.0<sup>§</sup> | 203.0 | 1094.1<sup>§</sup> | 4.3 | 3127.1<sup>#</sup> | 9.7 |
| G-CSF | 1.0 | 0.9 | 3581.1<sup>§</sup> | 138.0 | 246.6<sup>§</sup> | 27.4 | 35180.1<sup>#</sup> | 0.1 |
| GM-CSF | 36.0 | 15.5 | 555.1<sup>§</sup> | 19.0 | 56.5 | 5.8 | 1052.1<sup>#</sup> | 127.0 |
| CXCL10 | 0.5 | 0.0 | 18575.1<sup>§§</sup> | 808.1 | 29.1<sup>§</sup> | 10.5 | 23778.1<sup>#</sup> | 110.0 |
| CCL2 | 44.0 | 15.0 | 152.2<sup>§</sup> | 41.0 | 147.2<sup>§</sup> | 3.7 | 177.2 | 1.7 |
| CCL5 | 2.8 | 0.5 | 2615.1<sup>§§</sup> | 58.0 | 4.0 | 0.2 | 2797.2 | 68.0 |

Data are expressed as means of pg/ml ± SD obtained from three independent experiments. P values are calculated between: i) untreated and TNF-α - or IL-17A-treated groups, <sup>§</sup> $p < 0.05$ and <sup>§§</sup> $p < 0.01$ ; ii) TNF-α + IL-17A- versus TNF-α-treated groups, <sup>#</sup> $p < 0.05$ .

In the IL-36γ assay, we used a different Bio-Plex Pro™ assay kit. As reported in Table 2, treatment with IL-36γ alone up-regulated the secretion of CXCL1 and CXCL8. Consistently with previous data demonstrating the transcriptional induction by IL-36γ of CX3CL1, CXCL2, CXCL8, CCL2 and IL-6 mRNA in HDMECs [28], we found an up-regulation of the release of these proteins in HDMECs following IL-36γ treatment. An additive effect of IL-36γ and TNF-α could be observed for the chemokines CXCL5, CX3CL1, GM-CSF, CXCL1, CXCL2, CXCL8, CXCL10, CCL13, and for the IL-6 cytokine as compared to cells treated with TNF-α alone.

**Table 2.**
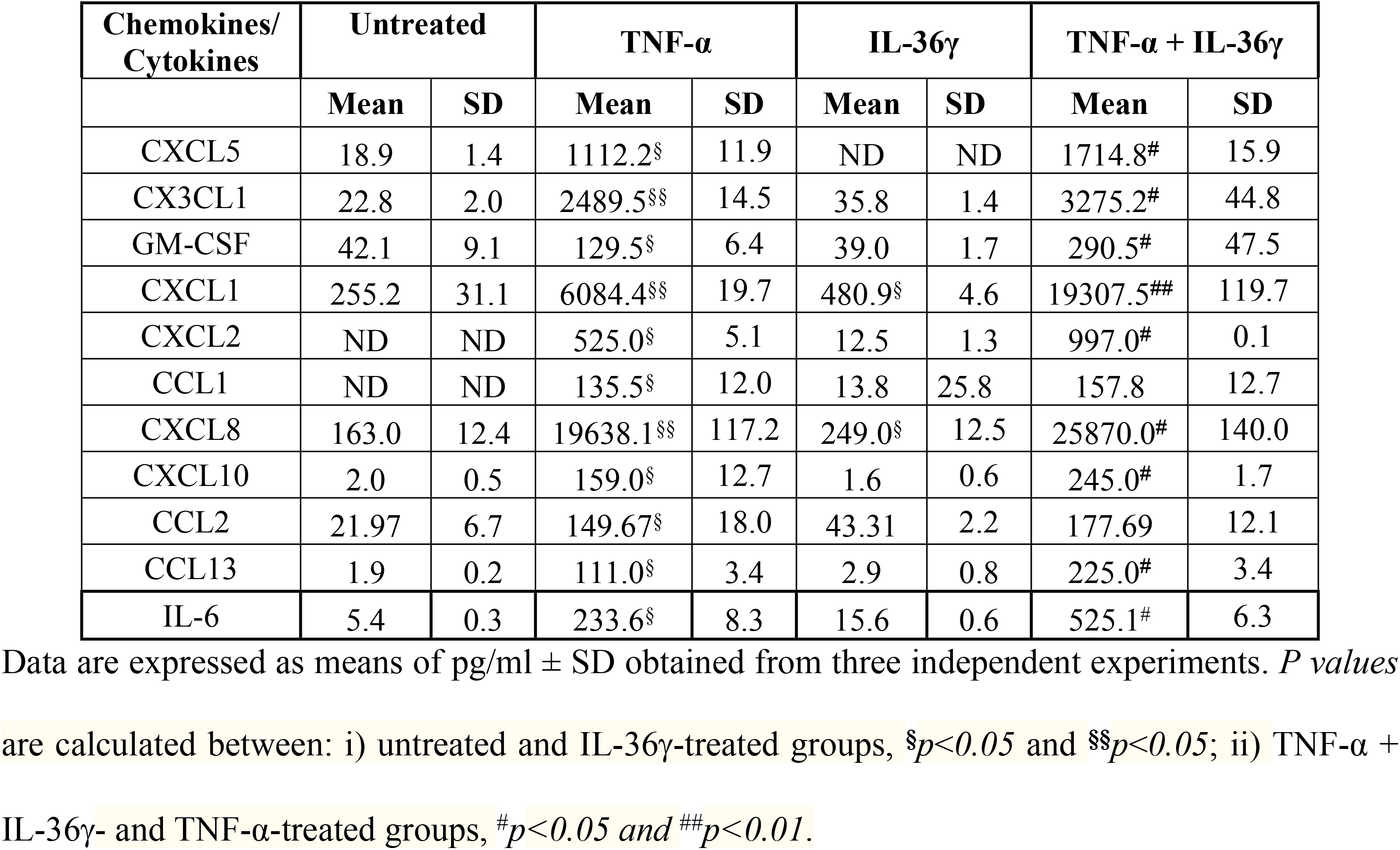
Bio-Plex 40-Plex chemokine analyses of supernatants from HDMECs treated with IL-36γ alone or in combination with TNF-α.

To validate these data, HDMECs were treated with IL-17A or IL-36γ, alone or in combination with TNF-α, and ELISA were used to measure the amounts of secreted inflammatory mediators selected from the Luminex assays. Analysis of secreted proteins confirmed most of the results obtained in these assays (Supplementary Table 1).

The pro-angiogenic role of IL-17A has been often ascribed to its ability to stimulate skin keratinocytes to release angiogenic factors, especially VEGF-A [31]. Thus, we analyzed VEGF-A protein secretion by human keratinocytes treated or not with IL-17A and combination of IL-17A and TNF-α. As shown in Fig 4A, IL-17A alone does not significantly induce the secretion of this angiogenic growth factor but it strongly synergizes with IFN-γ and TNF-α in stimulating its release by human keratinocytes.

**Fig 4.**
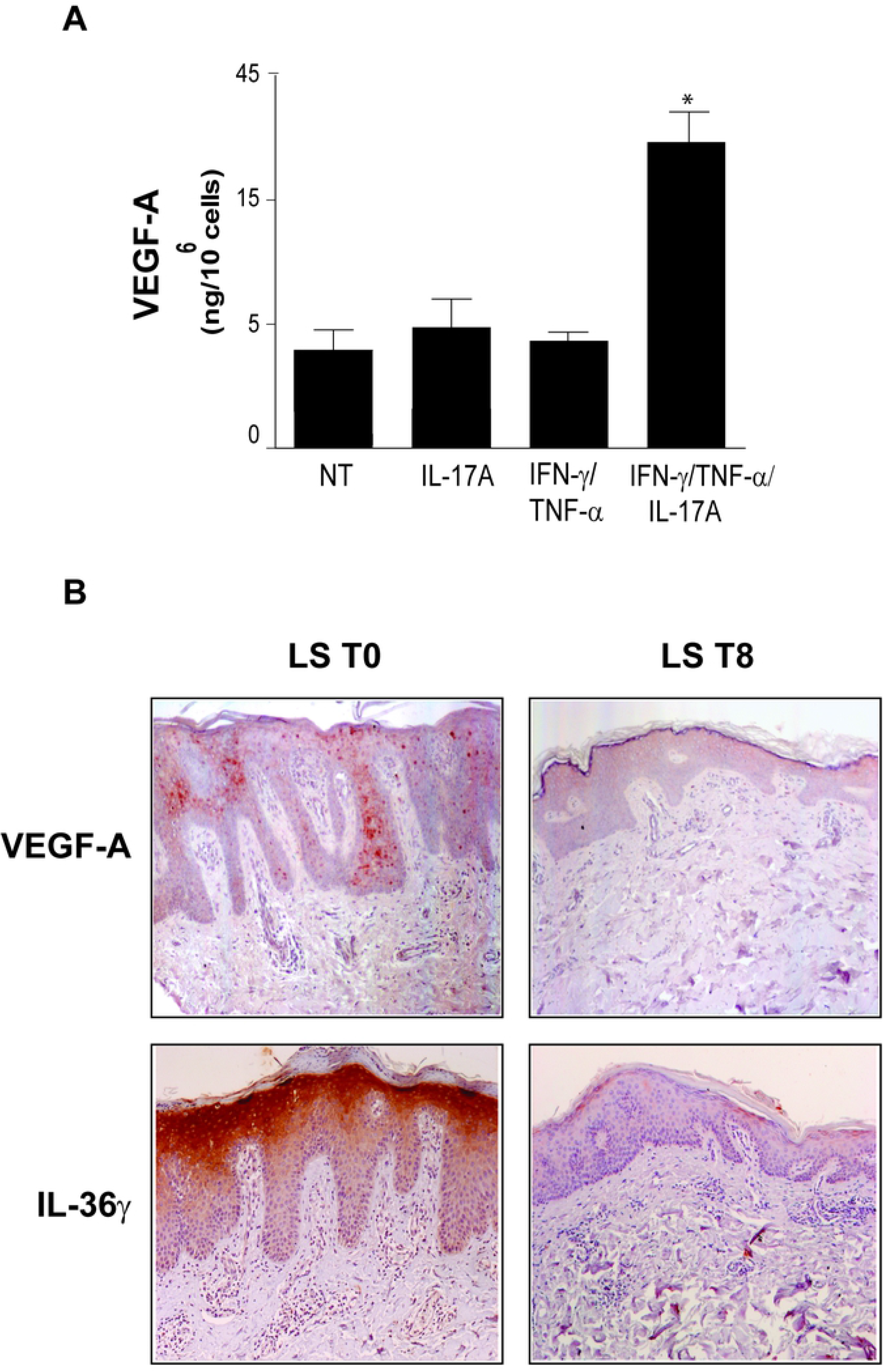
IL-17A in combination with TNF-α induces VEGF-A secretion both *in vitro* and *in vivo*. **A.** Supernatants from three different human keratinocyte strains were analysed for VEGF-A secretion by ELISA after treatment with IL-17A, alone or in combination with TNF-α and IFN-γ. NT= non treated. Results are presented as the mean of ng/10^6^ cells ±SD of the values obtained from the different strains in three independent experiments, \**p*<0.05 in respect to NT cells. **B.** VEGF-A or IL-36γ immunohistochemical staining (red) of patients’ lesional psoriatic skin (LS), before (T0) and after an eight-week treatment with Secukinumab (T8). Representative sections of skin specimens from three patients are shown (bars represent 100 μm).

There is evidence that VEGF-A is a primary angiogenic factor in psoriasis [32]. Serum levels of VEGF-A are higher in patients affected by psoriasis than in healthy controls, correlate with the Psoriasis Area and Severity Index (PASI) and diminish after treatment with psoralen plus ultraviolet-A (PUVA) or acitretin [33]. However, limited data are available about VEGF-A lowering after treatment of patients with anti-IL-17 antibodies. Thus, we analyzed *in vivo* VEGF-A expression after patient treatment with the anti-IL-17A antibody Secukinumab. As shown in Fig 4B, VEGF-A was strongly expressed in the suprabasal keratinocyte layer of the lesional skin and it was reduced after Secukinumab treatment for 8 weeks (Fig 4B). Consistently with previous reports [28], IL-36γ also accumulated in the upper layers of affected skin lesions, and its expression was drastically reduced by anti-IL-17A treatment.

To further clarify the role of VEGF-A in the IL-17/IL-36 axis and in the cross-talk between skin keratinocytes and HDMECs, we incubated HDMECs with culture medium conditioned by keratinocytes treated with IL-17A alone or in combination with TNF-α and analyzed proliferation and adhesion molecule expression in HDMECs. In parallel, we treated HDMECs with the IL-36 receptor antagonist (IL36RA) or with an inhibitor of VEGF-A signaling, the tyrosine kinase inhibitor Sunitinib, that blocks the activity of either VEGF or platelet-derived growth factor (PDGF) receptors as well as the signaling associated to CD117/c-kit [34]. We firstly verified the release of IL-36 isoforms and VEGF-A in the supernatants of untreated and cytokine-treated keratinocytes. Accordingly to data reported in Figures 1C and 4A, we found that both IL-36γ and VEGF-A were constitutively released and up-regulated upon IL-17A/TNF-α treatment, whereas IL-36γ was the cytokine mostly released by IL-17A-stimulated keratinocytes in their supernatants (data not shown). As shown in Fig 5A, stimulation with supernatants of untreated keratinocyte induced HDMEC proliferation compared to the EBM control, and cell proliferation was significantly reduced by both IL-36RA or Sunitinib. These results fit with the similar secreted amounts of IL-36γ and VEGF-A observed in the conditioned medium of untreated keratinocytes and thus with the angiogenic effect of these cytokines on HDMECs. Importantly, HDMEC treatment with the supernatant of IL-17A-stimulated keratinocytes further increased cell proliferation and this increment was blocked by IL-36RA but not by Sunitinib (Fig 5A), accordingly to the higher secretion of IL-36γ compared to VEGF-A in this condition. Therefore, blocking of VEGF-A signaling would result into an unnoticeable inhibition due to the contemporary presence of high amounts of an additional angiogenic factor such as IL-36γ. Unexpectedly, stimulation with both IL-17A and TNF-α-treated keratinocyte supernatants did not further induce HDMEC proliferation compared to untreated keratinocyte conditioned medium, but inhibition of either VEGF-A or IL-36γ was again effective in reducing cell proliferation (Fig. 5A). In fact, both VEGF-A and IL-36γ are likewise secreted by keratinocytes treated with a combination of TNF-α and IL-17A (Fig 1A, Fig 4A) and inhibition of either of them would be notable. In parallel to proliferation studies, FACS analyses of HDMECs treated with keratinocyte conditioned media showed that supernatants of IL-17A/TNF-α-treated keratinocytes slightly up-regulated ICAM-1 expression in HDMECs (Δmfi 101.5 compared to 61.6 of HDMECs grown in EBM medium, Fig 5B). Interestingly, IL-36RA reduced ICAM-1 membrane expression in HDMECs stimulated with supernatants of untreated or IL-17A- and IL-17A plus TNF-α-treated keratinocyte, whereas Sunitinib up-regulated ICAM-1 expression in all the three experimental conditions (Fig 5B). VCAM-1 expression by HDMECs was not influenced either by treatments with supernatants of cytokine-treated keratinocytes or by IL-36RA or Sunitinib inhibition (data not shown).

**Fig 5.**
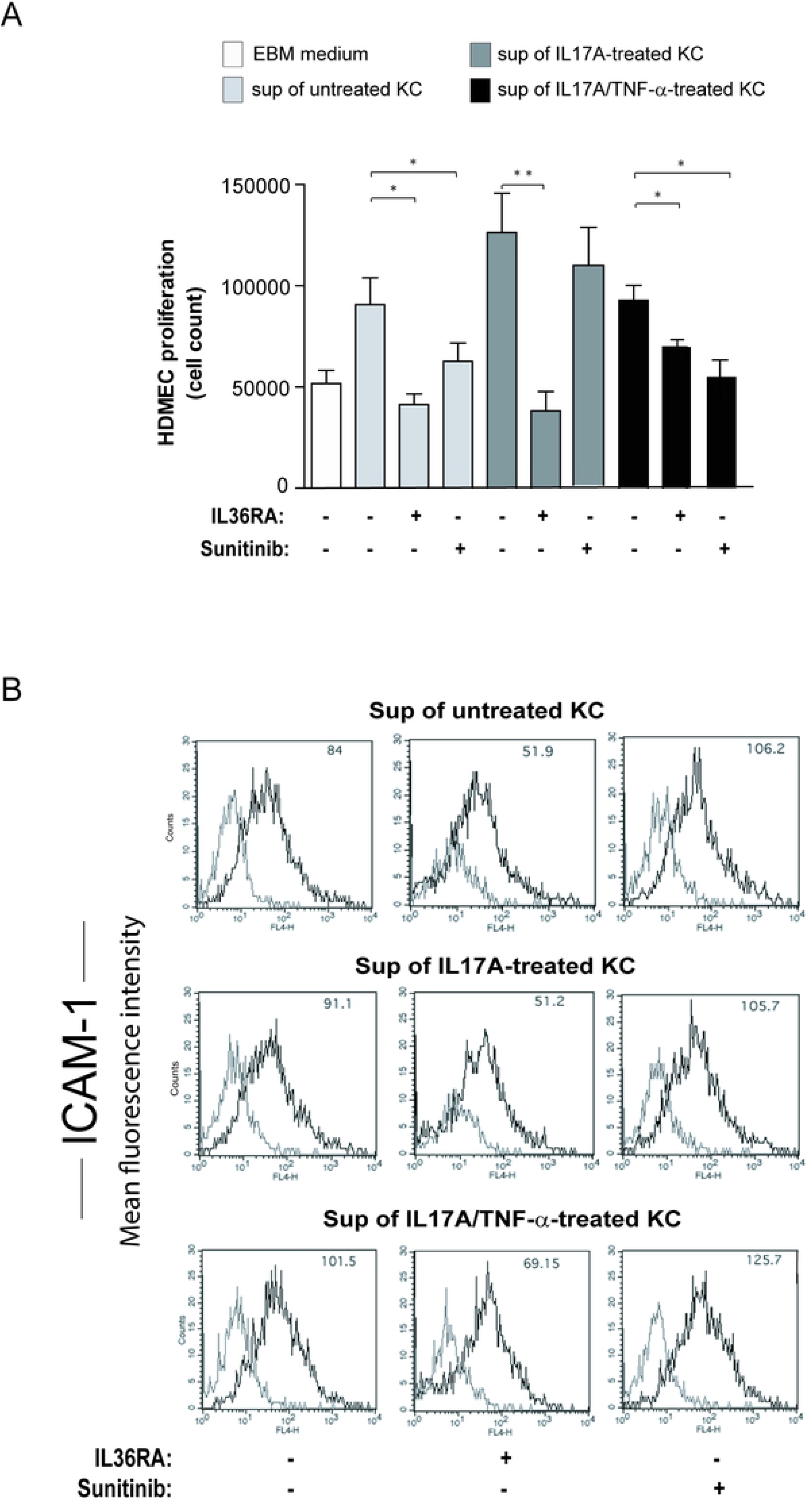
IL-36R activation mediates the angiogenic effects of IL-17A. **A.** Cell proliferation of HDMECs treated for 24 hours with the basal medium (EBM), supernatant of untreated, IL-17A- or of IL-17A/TNF-α-treated keratinocytes (KC) was evaluated by viable cell counts. IL-36RA or Sunitinib were added during the assay, as indicated. Data are shown as mean values of cell counts of three samples ± SD. Experiments were repeated at least three times with similar results; \**p*≤0.05, ** *p*≤0.01. **B.** ICAM-1 expression was evaluated by flow cytometry analysis on HDMECs stimulated for 24 hours with the supernatant of untreated, IL-17A- or IL-17A/TNF-α-treated KC in the presence or not of IL-36RA or Sunitinib. Results are shown as Δmfi. A representative experiment out of three performed is shown. Δmfi of ICAM-1 expression in HDMECs grown in EBM was 61.6.

## Discussion

It is well known that both IL-17A and IL-36γ activate pathogenic pathways in different cell types in psoriatic skin, especially in resident skin cells such as keratinocytes and dermal EC. While much is known about the direct impact of IL-17A and IL-36γ on these skin cell types, no studies about their indirect inflammatory effects on EC mediated by keratinocytes have been reported so far. In this work, we highlight the strong IL-17A- and IL-36γ-dependent interplay between keratinocytes and ECs in psoriatic condition that leads to the establishment of a cytokine network responsible for the development and maintenance of the inflamed state. In particular, we clarify the action of IL-17A in the cross-talk between skin keratinocytes and ECs by investigating the involvement of IL-36γ and VEGF-A, both abundantly and constitutively produced by keratinocytes and augmented after cytokine stimulation. Our study indicates that IL-17A promotes EC proliferation *in vitro* both directly and indirectly through induction of IL-36γ release by keratinocytes. Moreover, IL-17A synergizes with TNF-α in inducing both IL-36γ and VEGF-A in human keratinocytes. Therefore, when HDMECs are stimulated with the supernatant of IL-17A-treated keratinocytes, IL-36γ, but not VEGF-A, is the more abundantly released cytokine in the medium and is likely responsible for the observed increase of HDMEC proliferation. Consistently, inhibition of IL-36γ by IL-36RA, but not of VEGF-A by Sunitinib, reduced EC proliferation. Our results support the idea that IL-36 cytokines and, in particular, IL-36γ are important angiogenic mediators of IL-17A. On the other hand, stimulation of HDMEC proliferation with a supernatant of IL-17A- and TNF-α-treated keratinocytes, where equally augmented amount of IL-36γ and VEGF-A are present, could be inhibited by either IL36RA or Sunitinib. Interestingly, treatment with supernatants of IL-17A- and TNF-α-treated keratinocytes did not further enhance HDMEC proliferation in respect to stimulation with supernatants of untreated keratinocytes. Therefore, we can speculate that the IL-17A/TNF-α combination induces the release by keratinocytes of additional mediators able to counteract the proliferative action of IL-36γ and/or VEGF-A on HDMECs. It is evident that the angiogenic effect of IL-17A is precisely tuned by different proteins present in the inflamed skin. Our data appear in contrast with those by Bridgewood et al. [35] where IL-36-stimulated EC tubulogenesis was significantly impaired by either an anti-VEGF-A neutralizing antibody or by Sunitinib. In our hands, IL-36γ did not induce VEGF-A production by either keratinocytes or HDMECs *in vitro* (data not shown). However, there is the possibility that other factors induced by IL-36-treatment on ECs could indirectly stimulate VEGF-A release that in turn stimulated tubule formation.

Concerning the angiogenic role of IL-17A in psoriasis, it is important to emphasize the direct effect of this cytokine on the proliferation of IL-17R-bearing ECs, as previously reported [14,17] and confirmed in this study. At the molecular level, we observe that IL-17A supports EC proliferation by inducing activation of either NF-κB-, ERK1/2- or STAT3-dependent pathways, known to be implicated in proliferation and survival of several cell types. Through activation of IL-36R, also IL-36γ directly induces EC proliferation, even though at a lower extent than IL-17A, possibly activating NF-κB and ERK1/2 signaling. The stronger activation of ERK1/2 and STAT3 pathways by IL-17A compared to that obtained with IL-36γ could explain the more pronounced mitogenic effect of IL-17A [26,36,37].

Other than having a mitogenic effect, we demonstrate that IL-17A counteracts the anti-proliferative effect of TNF-α, potentially activating the pro-survival ERK1/2 pathway. Indeed, IL-17A could protect keratinocytes from the pro-apoptotic effect of TNF-α, but a more detailed analysis of the molecular mechanisms underlying this process should be performed. The identification of these mechanisms could also explain why IL-36γ, even if able to stimulate the ERK1/2-dependent pathway, did not counteract anti-proliferative effects of TNF-α as efficiently as IL-17A. Our analysis of the inflammatory responses in dermal ECs reveals that keratinocyte-derived IL-36γ contributes to the expression of ICAM-1, necessary to leukocyte binding to vessel and extravasion into the tissue, as demonstrated by inhibiting IL-36 action on HDMECs treated with supernatants derived from keratinocytes. ICAM-1 is particularly inhibited when HDMECs are stimulated with supernatants from keratinocytes treated with IL-17A plus TNF-α, a combination of cytokines particularly efficacious in up-regulating IL-36 cytokines, and in particular IL-36γ. However, considering that IL-36γ alone does not directly induce ICAM-1 expression by HDMECs, it is possible that other IL-36 isoforms, such as IL-36β, could be responsible for the observed ICAM-1 expression. Of note, blocking of VEGF-A in keratinocyte-derived supernatants by Sunitinib treatment results into up-regulation of membrane ICAM-1. Similar results have been previously obtained with different VEGF-A inhibitors [38]. Induction of ICAM-1 expression by VEGF-A inhibition is an emerging aspect that supports the association of immunotherapy and anti-angiogenic therapy in cancer treatment. In fact, blocking angiogenesis could make the tumor more accessible and vulnerable to the immune system [39]. In our experimental model, the combination of Sunitinib with pro-inflammatory cytokines present in the cytokine-treated keratinocyte medium or expressed by ECs exposed to the keratinocyte supernatants, could result into ICAM-1 up-regulation as well. A recent paper by van Hooren and colleagues [40] demonstrated the up-regulation of ICAM-1 expression on tumor ECs treated with Sunitinib and an agonistic anti-CD40 monoclonal antibody. CD40 is a member of the TNF super family and, in immune cells, its expression is induced by several pro-inflammatory cytokines such as IL-36γ itself. These pro-inflammatory effects of VEGF inhibition could be responsible for the limited results of anti-angiogenic treatment of psoriasis, where high amounts of VEGF-A as well as prominent angiogenesis is present [41].

Despite what previously reported [11], we clearly demonstrate that IL-17A alone has a direct pro-inflammatory effect on ECs, by inducing cytokines and chemokines, such as IL-6, CXCL8, G-CSF, CXCL10 and CCL2. In addition, IL-17A synergized with TNF-α on ECs, as indicated in other studies [13], either through induction of membrane proteins, such as ICAM-1, or by secretion of inflammatory soluble mediators.

Our data support the hypothesis that targeting IL-17A should result in an improvement of the EC damage observed in psoriasis patients. This chronic microvascular damaging would lead through time to the cardiovascular co-morbidities recurrently associated to psoriasis. Clinical trials indicate that treatment with biological drugs, such as Secukinumab, that target the IL-17A signaling pathway, markedly improves disease outcome. These IL-17-targeting drugs are generally well tolerated and constitute a good alternative to other biological compounds that target TNF-α. Anti-TNF-α treatment with Infliximab of psoriasis patients significantly reduce the levels of VEGF-A [42]. Similar data are not fully available for the IL-17-targeting compounds. As for Secukinumab, the 52-week, randomized, double-blind, placebo-controlled, exploratory trial CARIMA showed a tendency of psoriasis patients in ameliorating endothelial functions measured by flow-mediated dilation [43]. On the other hand, in mouse models of psoriasis, vascular inflammation, evaluated through the measure of circulating inflammatory cytokines and chemokines, and vascular dysfunction, analyzed *ex-vivo* by vascular responsiveness to vasodilators, were correlated with the severity of skin lesions and levels of IL-17A. In a model of moderate to severe form of psoriasis with a late onset, anti-IL-17A treatment have beneficial effects on both vascular inflammation and dysfunction [44]. Our results are consisting with these findings and further indicate that Secukinumab treatment reduces VEGF-A together with IL-36γ presence in the psoriatic skin. This double reduction could effectively counteract a possible induction of inflammatory membrane molecules due to VEGF-A diminution. In the future, for a better management of psoriasis cardiovascular co-morbidities, the association of anti-IL-17A therapy with an anti-IL-36γ treatment should be taken into account.

## Acknowledgments

Authors would like to thank Novartis Farma Italy for participating in funding this preclinical project. This work was supported by grants from the Italian Ministry of Health “Ricerca Corrente”. The technical support of the Complex Protein Mixture (CPM) analysis facility at ISS is kindly acknowledged.

